# State-Dependent Regulatory Compression: Chromatin Geometry Gates Information Flow in Hematopoiesis

**DOI:** 10.64898/2025.12.31.697240

**Authors:** Roberto Navarro Quiroz, Elkin Navarro Quiroz

**Affiliations:** Center for Research in Critical Dynamics, Barranquilla, Colombia; Universidade Estadual Paulista (UNESP), Instituto de Química, Araraquara, Brazil; Universidad Simón Bolívar, Centro de Investigaciones en Ciencias de la Vida (CICV), Barranquilla, Colombia

## Abstract

Geometric constraints in chromatin-transcription space—regions of low occupancy termed “forbidden zones”— have been interpreted as signatures of regulatory dissociation in progenitor cells. We previously falsified this interpretation: progenitors exhibit higher, not lower, mutual information between chromatin accessibility and transcription. Here we address the consequent question: what organizational principle governs coupling under geometric constraint? Using human bone marrow multiome data (GSE194122; N=13 donors, 69,249 cells), we operationalize the Law of State-Dependent Regulatory Compression (LCR-DE) through three metrics: Gate Occupancy (GO), Coupling Efficiency (CE), and Control Selectivity (CS). Progenitors maintain conserved geometric boundaries (GO: Wilcoxon p=0.94) while exhibiting elevated coupling efficiency (CE: median 0.058 vs 0.019; p=9.77× 10^−3^). The lower CS in progenitors reflects distributed regulatory redundancy across multilineage programs, not absence of control, while differentiated cells exhibit crystallized, pathway-specific channeling. CE persists after cell cycle residualization (p=0.64), confirming biological rather than proliferative origin. These findings establish regulatory compression—informational channeling through selective pathways under geometric constraint—as an organizational principle of hematopoietic differentiation.

## Introduction

Single-cell multiome profiling has revealed that the joint distribution of chromatin accessibility and gene expression occupies a geometrically constrained manifold (Cao et al., 2018; Ma et al., 2020). The high-ATAC/low-RNA quadrant—the “forbidden zone”—is systematically underpopulated, particularly in progenitor populations. This observation invited the interpretation that developmental plasticity operates through informational decoupling: accessible chromatin that does not translate to transcription (Ma et al., 2020; Buenrostro et al., 2018).

This interpretation conflates two distinct properties. Geometric constraint describes *where* cells can reside in bivariate space; informational coupling describes *how precisely* one variable predicts another within that space. A distribution can be geometrically restricted while exhibiting maximal mutual information within that region— compression of the state space typically increases, not decreases, information density per unit volume.

We recently addressed this gap directly by measuring mutual information (MI) between chromatin accessibility and transcription in progenitor versus differentiated cells (Navarro Quiroz and Navarro Quiroz, 2025). The result falsified the dissociation hypothesis: progenitors exhibit higher, not lower, MI. Geometric constraint coexists with elevated regulatory coupling.

This negative result opens a constructive question that the present work addresses: *if the forbidden zone does not represent entropic dissociation, what organizational principle does it implement?*

Here we propose and test the **Law of State-Dependent Regulatory Compression (LCR-DE)**: geometric constraints implement informational compression by channeling regulatory coupling through selective, state-dependent pathways. Under this framework, the forbidden zone functions as a gate that enhances signal-to-noise ratio, not as a barrier that blocks information flow. The conceptual foundation: geometric constraint ≠ entropic decoupling.

We operationalize LCR-DE through three complementary metrics. **Gate Occupancy (GO)** quantifies the geometric boundary—the fraction of cells traversing the forbidden zone. **Coupling Efficiency (CE)** measures the informational capacity of the chromatin-to-transcription channel after removing technical and proliferative confounders. **Control Selectivity (CS)** tests whether coupling operates through specific regulatory features or through generic genome-wide accessibility—distinguishing crystallized from distributed control architectures.

## Results

### Cohort composition and analytical validity

The GSE194122 dataset comprises 13 donors across 4 collection sites, totaling 69,249 cells after quality control (Table S1). Progenitor abundance varied substantially: from 1,421 cells (donor8_site4) to 94 cells (donor7_site3). One donor (donor5_site2) contained no annotated progenitor cells. For CE and CS analyses requiring robust MI estimation, we imposed a minimum threshold of 200 progenitor cells, yielding N=10 donors with valid paired measurements. GO analysis retained N=12 donors. Site-stratified analysis confirmed pattern reproducibility across independent collection centers (Supplementary Figure 1).

### Geometric gating is conserved across developmental states

Gate Occupancy quantifies the fraction of cells residing in the forbidden zone, defined by thresholds 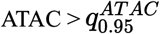 and 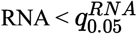 .Critically, these quantiles were computed per donor using **all cells** (Progenitor +Differentiated+ Others) *before* state subsetting, establishing a fixed geometric reference frame not auto normalized by state composition (Table S2). Donor-specific thresholds ranged from 9.35 to 10.12 loglp units for ATAC and 5.62 to 6.99 logIp units for RNA.

Figure 1 presents GO across developmental states. The null result is striking: no significant difference between progenitors and differentiated cells (Wilcoxon signed-rank p = 0.94, N=12). Both populations exhibit low GO (median < 0.001), indicating that the geometric boundary is similarly restrictive across developmental states.

**Figure 1.**
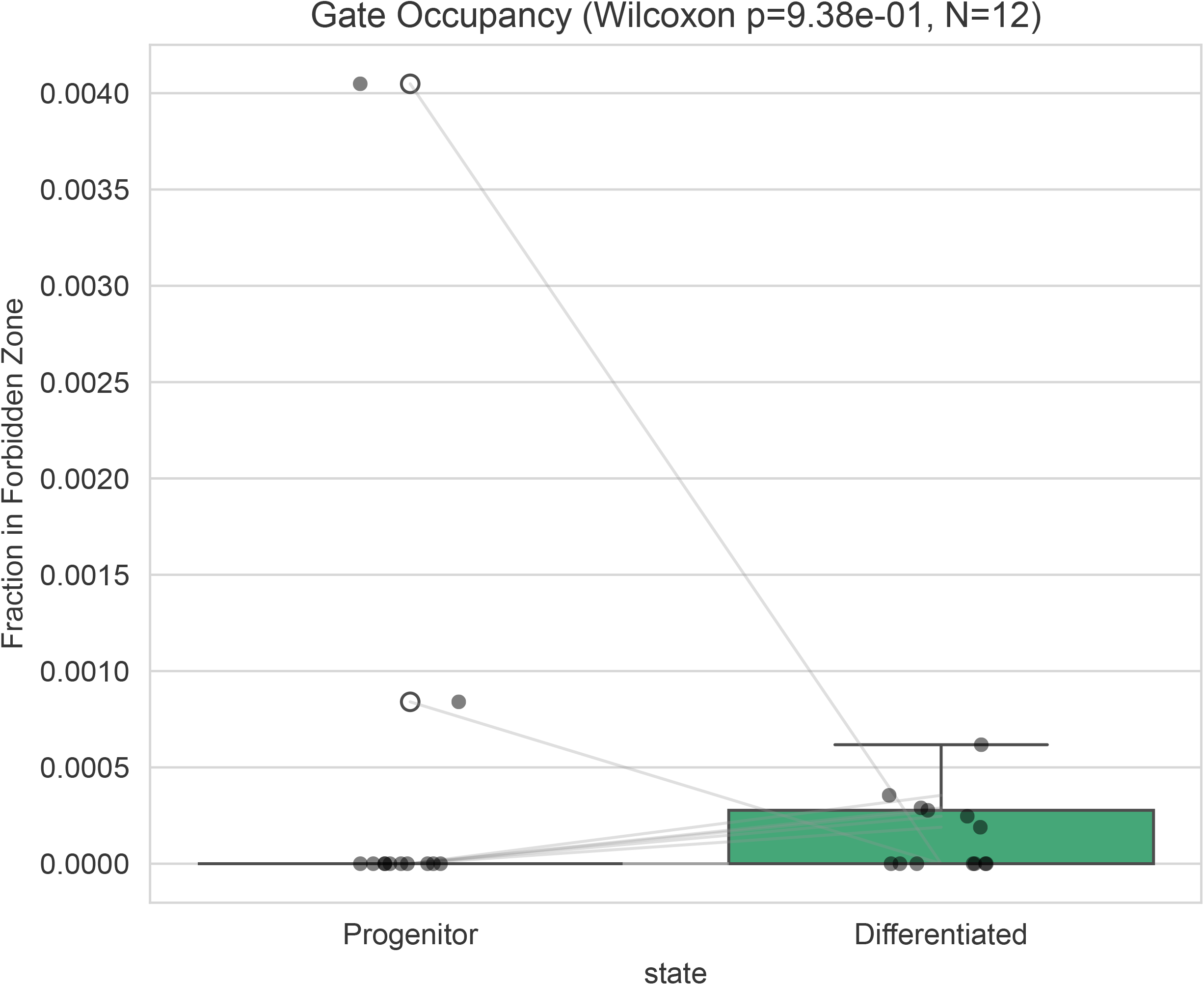
Gate Occupancy (GO): Geometric boundaries are conserved across developmental states. Box plots comparing GO (fraction of cells in forbidden zone) between Progenitor (gray) and Differentiated (green) populations. Paired donor observations connected by lines. Wilcoxon signed-rank p = 0.94, N=12 donors. Both states exhibit low GO (median< 0.001), indicating the geometric gate is preserved throughout hematopoiesis. The null result demonstrates that what changes upon differentiation is not the boundary, but information flow through it. Thresholds computed per donor using all cells before state subsetting (Table S2).

This null result is informative, not empty. The forbidden zone is a conserved organizational feature—it does not relax upon differentiation. What changes between states is not whether the boundary exists, but how information flows through the boundary. The gate structure is preserved; the channel properties differ.

### Coupling efficiency is elevated in progenitors

If geometric constraint implied informational dissociation, progenitors should exhibit reduced coupling efficiency. The data reject this prediction.

Coupling Efficiency (CE) was estimated as KSG mutual information (k=10 neighbors) on depth-residualized, copula-transformed accessibility and expression values. The input variables were:

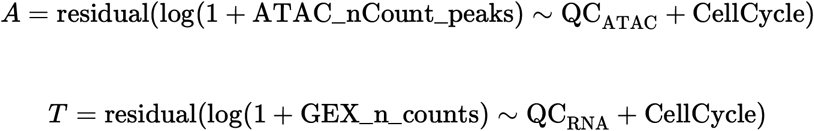

where QC terms include fragment counts, nucleosome signal, and mitochondrial fraction, and CellCycle includes S-phase and G2M scores.

CE is significantly higher in progenitors than differentiated cells (Figure 2; Wilcoxon stat= 3.0, p= 9.77 × 10^− 3^ N=10). Median progenitor CE (0.058) exceeds median differentiated CE (0.019) by 3.1-fold. The effect is consistent: 9 of 10 donors show CE_prog > CE_diff Comparison with permutation null models (C2 control) confirms that empirical CE substantially exceeds chance—null CE clusters near zero while empirical progenitor CE reaches 0.05-0.18 (Table S5).

**Figure 2.**
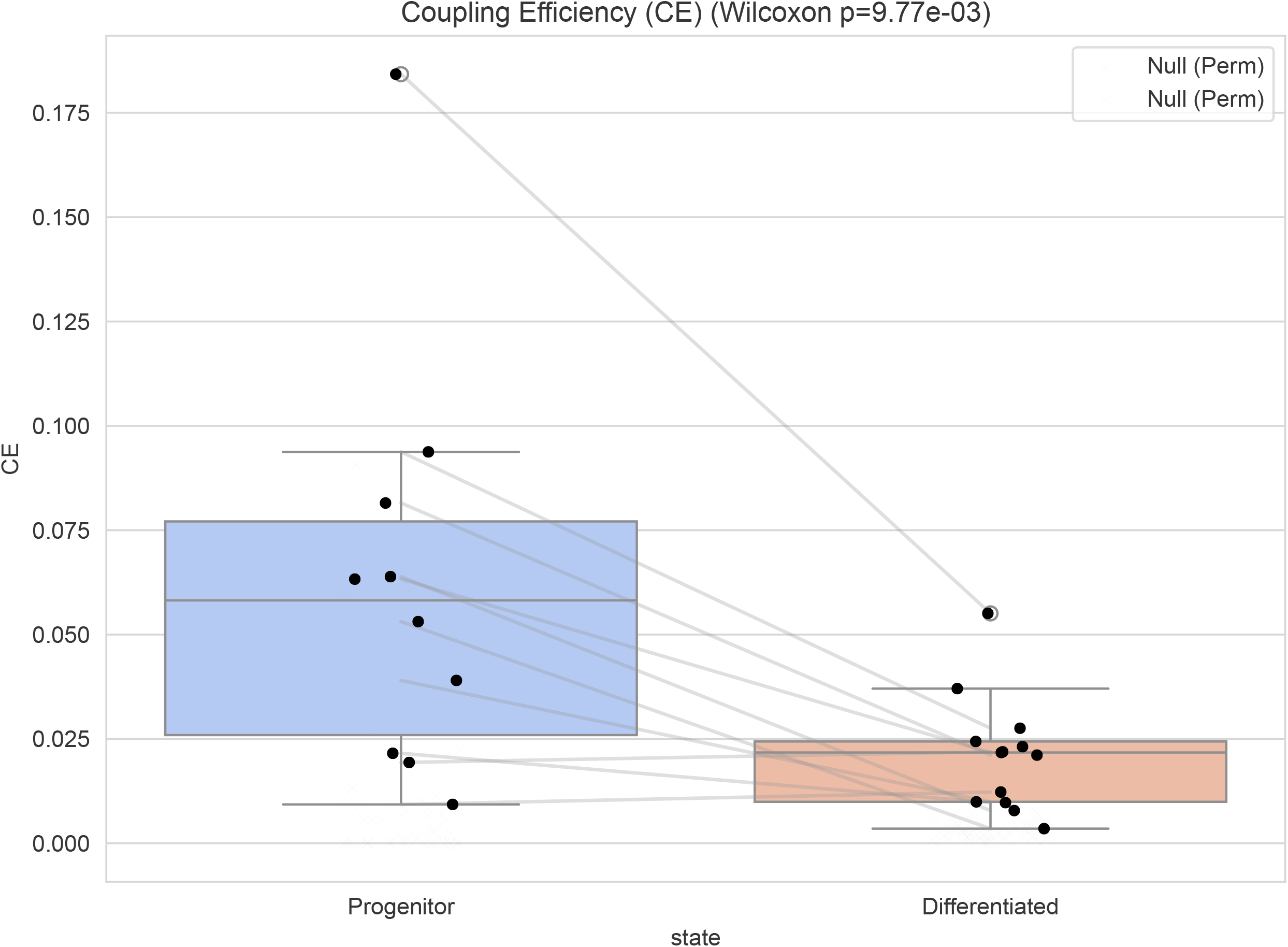
Coupling Efficiency (CE): Progenitors exhibit elevated chromatin-transcription information capacity. Box plots comparing CE (KSG mutual information on depth- and cell cycle-residualized, copula-transformed values) between states. Blue/coral boxes: empirical estimates; x markers: permutation null (C2 control). Paired observations connected. Progenitors exhibit significantly higher CE (median 0.058 vs 0.019; Wilcoxon p = 9.77 × 10^− 3^, N=lO). Null CE clusters near zero, confirming genuine dependence. The 20-fold range in progenitor CE (0.009-0.184) reflects biological heterogeneity in regulatory organization.

#### State-label shuffling within donors abolishes CE/CS differences (data not shown)

This C4 control confirms that observed patterns are genuinely state-specific.

### Control Selectivity reveals distinct regulatory architectures

Control Selectivity (CS) measures whether coupling operates through specific regulatory features or diffuse genome-wide accessibility:

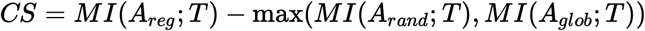

where *A*_*reg*_ is PCI of the top 2000 differentially accessible peaks between states, *A*_*rand*_ is PCI of accessibility-matched random peaks, and *A*_*glob*_ is global accessibility.

Figure 3 presents CS by developmental state. Differentiated cells exhibit significantly higher CS than progenitors (Wilcoxon p = 2.73×10^−2^, N=10). This pattern requires careful interpretation that elevates rather than weakens the framework.

**Figure 3.**
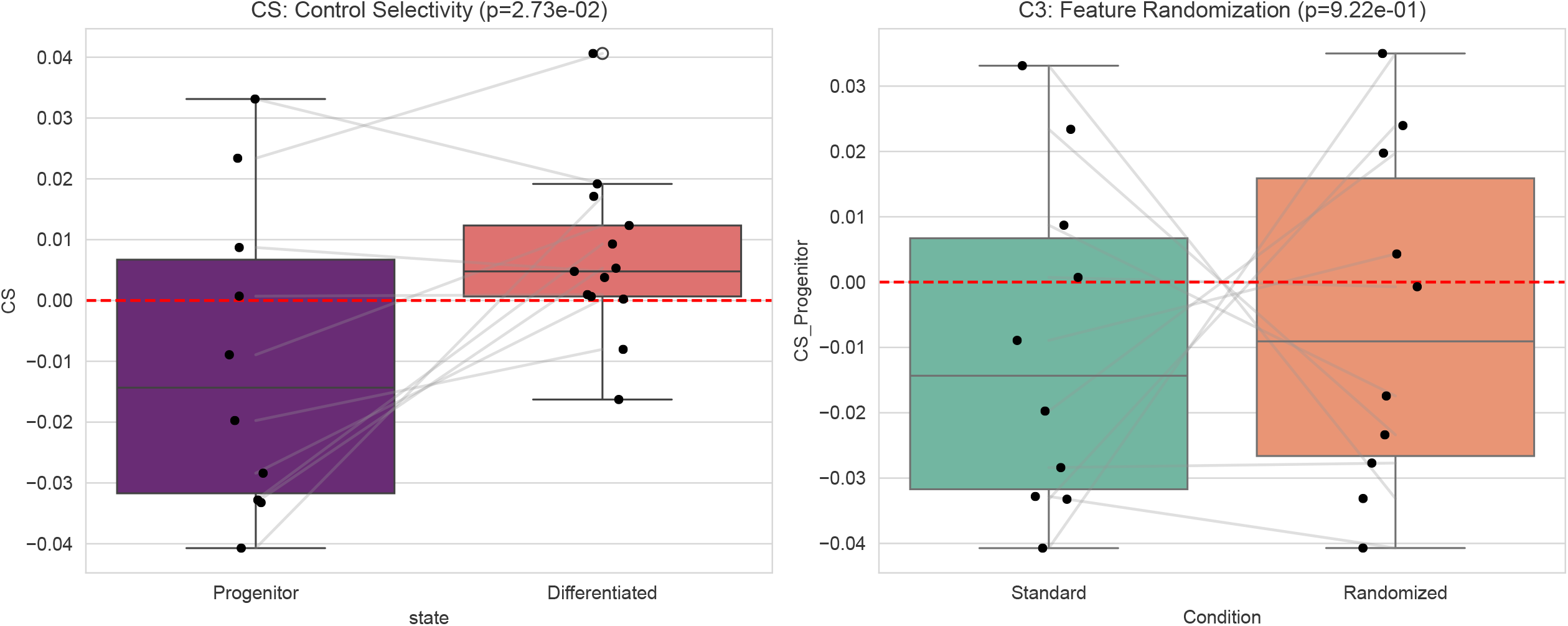
Control Selectivity (CS): Distinct regulatory architectures between developmental states. Left panel: CS by state. Differentiated cells exhibit higher CS (Wilcoxon p = 2.73×10^−2,^ N=lO), reflecting crystallized, pathway-specific regulatory channeling. Lower, variable progenitor CS reflects **distributed regulatory redundancy—a** holographic architecture where control spreads across multilineage programs rather than concentrating in one pathway. This is the informational signature of multipotency, not absence of selectivity. Red dashed line: CS = 0. Right panel (C3 control): Feature randomization abolishes pattern (p = 0.92), confirming signal requires regulatory feature identity.

In **differentiated cells**, the regulatory landscape has crystallized. A defined set of lineage-specific enhancers drives transcription; coupling flows through these specific features, yielding positive CS. The differentiated state has resolved its regulatory architecture into discrete, pathway-specific channels.

In **progenitors**, CS is lower and more variable, including negative values. This reflects **distributed regulatory redundancy—a** holographic architecture where control is not concentrated in any single pathway but spread across multilineage programs maintained simultaneously. The binary *F*_*reg*_ definition (progenitor vs differentiated contrast) systematically underestimates this distributed architecture because it captures only one axis of a multidimensional regulatory space.

The lower progenitor CS does not indicate “noise” or “absence of selectivity.” It indicates **delocalized control** -regulatory information distributed across multiple potential lineage programs rather than concentrated in one crystallized pathway. This is the informational signature of multipotency: high total coupling (CE) with distributed rather than focal channeling (CS).

The C3 control (Figure 3, right panel) validates that CS depends on regulatory feature identity: randomizing features abolishes the progenitor-differentiated CS distinction (p = 0.92).

### Cell cycle does not account for coupling efficiency

A potential confound is that progenitors exhibit higher proliferative activity, which could drive apparent chromatin-transcription coupling through cell cycle-dependent gene expression. Figure 4 presents CE before and after residualizing cell cycle scores (S-phase and G2M).

**Figure 4.**
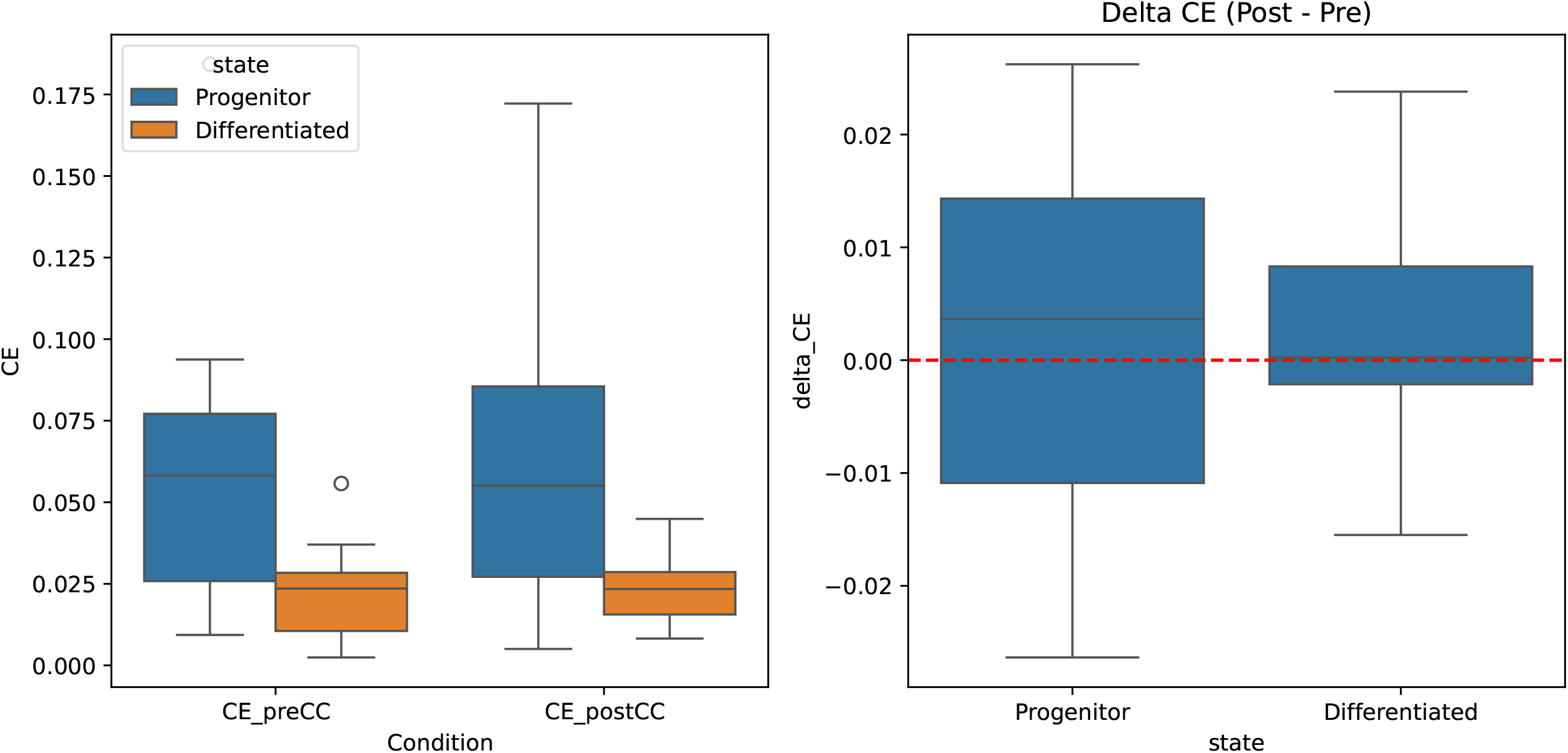
Cell Cycle Control: Coupling efficiency persists after proliferative residualization. Left panel: CE before (preCC) and after (postCC) cell cycle score residualization, stratified by state. Progenitor (blue) > Differentiated (orange) pattern maintained in both conditions. Right panel: ΔCE (postCC - preCC) by state. Distributions center near zero with no systematic shift (Wilcoxon p = 0.64 for pre vs post). Elevated progenitor CE reflects regulatory organization, not proliferative state.

CE is not significantly altered by cell cycle residualization (Wilcoxon p = 0.64, N=23 observations). The progenitor-differentiated CE difference persists: median progenitor ΔCE remains approximately 3-fold higher than differentiated CE in both conditions. The right panel confirms that ΔCE (post minus pre cell cycle correction) centers near zero in both states, with no systematic shift.

This control establishes that elevated progenitor CE reflects regulatory organization, not proliferative state. The chromatin-transcription channel operates independently of cell cycle phase.

### Hostile controls summary

All metrics were subjected to hostile controls designed to falsify artifactual interpretations (Table S6):

#### Cl (Depth + Cell Cycle Residualization):)

All metrics computed on residuals from within-donor regression against QC metrics and cell cycle scores. Effects persist.

#### C2 (Block Permutation)

Transcription values permuted within donor × state blocks (N=5 independent pe1mutations). Null MI distributions center near zero; empirical CE exceeds null by 5-50x (Table S5).

#### C3 (Feature Randomization)

CS computed using random (not accessibility-matched) features yields no systematic progenitor-differentiated difference (p = 0.92). Signal requires regulatory feature specificity.

#### C4 (Label Shuffle)

State-label shuffling within donors abolishes CE/CS differences (data not shown). Pattern is genuinely state-dependent.

#### CS (Threshold Sensitivity)

GO computed across quantile grids (*q*_*90*_*-q*_*99*_ for ATAC, qo1*-q10*for RNA). Pattern robust to threshold choice.

#### Site Robustness

CE and CS patterns reproduce across all four collection sites (Supplementary Figure 1, Table S10). No site shows reversed pattern direction.

All controls pass. The observed patterns—conserved geometric boundaries, elevated progenitor coupling efficiency, state-dependent selectivity architecture—reflect biological organization.

## Discussion

### Geometry implements compression, not dissociation

The central finding is that geometric constraints in chromatin-transcription space do not indicate informational decoupling. The forbidden zone is not a wall blocking signal propagation; it is a gate channeling information flow.

In communication theory, compression reduces bandwidth requirements while preserving signal fidelity (Cover and Thomas, 2006). A compressed channel appears geometrically restricted—fewer states accessible—while maintaining or enhancing effective information transfer. The chromatin-transcription channel in progenitors exhibits exactly this signature: geometric restriction (conserved GO) with elevated mutual information (higher CE).

This resolves the apparent paradox of chromatin priming. Progenitor cells maintain accessible chromatin at lineage-specific loci without maximal transcription. Previous interpretations viewed this as “wasteful” or “decoupled.” Under LCR-DE, primed chromatin is precisely functional: it maintains regulatory channels in a ready state, enabling rapid, high-fidelity transcriptional responses to differentiation signals.

### Distributed versus crystallized control

The CS metric reveals distinct regulatory architectures between developmental states. Differentiated cells exhibit crystallized control—information flows through defined pathway-specific channels, yielding positive CS. Progenitors exhibit distributed control-information spreads across multiple regulatory programs simultaneously, yielding lower or variable CS.

This distinction has physical content. A holographic system stores information distributively; damage to any single region does not catastrophically disrupt function. A focal system stores information locally; disruption to the critical pathway eliminates function. Progenitors appear holographic; differentiated cells appear focal.

The lower progenitor CS is not a weakness of the framework—it is a prediction confirmed. Multipotent cells maintaining multiple lineage options simultaneously should exhibit distributed rather than concentrated regulatory coupling. The *F*_*reg*_ definition, based on binary state contrast, necessarily underestimates this multilineage architecture. Future work should define lineage-specific feature sets to decompose progenitor CS into its constituent pathways.

### Relation to rate-distortion theory

The LCR-DE framework connects to rate-distortion theory in information theory (Cover and Thomas, 2006). Given a constraint on channel capacity (geometric restriction), optimal encoding minimizes distortion while respecting bandwidth limits. The progenitor state may approach this optimum: geometric boundaries define the available “bandwidth,” and elevated CE indicates efficient use of that bandwidth.

The differentiated state relaxes this constraint differently — not by expanding geometry, but by simplifying the encoding. Crystallized CS indicates that fewer regulatory degrees of freedom are active, reducing the complexity of the chromatin-transcription mapping while maintaining functional output.

### Explicit relation to Paper 1

This work extends, rather than repeats, our previous study. Paper 1 established a falsification: geometric constraint does not imply informational dissociation. Progenitors show higher, not lower, MI.

Paper 2 addresses the constructive question: given that constraint coexists with coupling, *how* is this achieved? The answer is state-dependent compression with distinct control architectures. GO documents preserved geometry. CE documents elevated information. CS documents the crystallized-versus-distributed architecture of that information flow. Cell cycle controls confirm biological rather than proliferative origin.

### Implications

#### For stem cell biology

The primed chromatin state implements distributed regulatory readiness, not entropic noise. Interventions that globally modify chromatin accessibility may disrupt this distributed architecture and paradoxically reduce regulatory precision.

#### For disease

Pathological stemness (leukemic stem cells) may involve disruption of the distributed control architecture—either inappropriate crystallization (loss of multipotency) or loss of geometric boundaries (uncontrolled accessibility). These represent distinct failure modes with different therapeutic implications.

#### For multiome methodology

Geometric features (forbidden zones) and informational features (CE, CS) are distinct properties requiring separate quantification. Visual inspection of scatter plots cannot distinguish dissociation from compression. Studies claiming “decoupling” should validate with information-theoretic metrics.

#### Limitations

The effective sample size for paired analysis is N=10 donors; larger cohorts would refine effect size estimates. The *F*_*reg*_ definition uses binary state contrast, likely underestimating multilineage progenitor selectivity. Cross-sectional design provides snapshots, not dynamics. Generalization beyond hematopoiesis requires explicit testing.

## Materials and Methods

### Data source

GSE194122 (Luecken et al., 2022): 10x Genomics Multiome from human bone marrow. After QC filtering, 69,249 cells from 13 donors across 4 sites.

### Gate Occupancy (GO)

Forbidden zones defined by 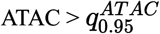 and 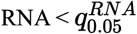. **Quantiles computed per donor using ALL cells (before state splitting) to avoid auto-no1malization** (Table S2). GO= fraction of cells in state falling within fixed boundaries.

### Coupling Efficiency (CE)

KSG mutual information *(k* = 10; Kraskov et al., 2004) on copula-transformed ranks of residualized values:

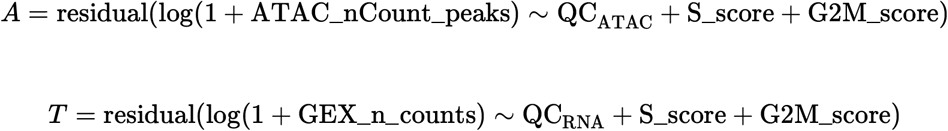

QC_ATAC: total fragments, nucleosome signal, TSS enrichment. QC_RNA: total counts, detected features, mitochondrial fraction. Cell cycle scores computed via scanpy (Tirosh et al., 2016). Subsampling: N=2000 cells, R=20 replicates.

### Control Selectivity (CS)

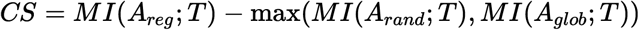

*F*_*reg*_:Top 2000 differentially accessible peaks (progenitor vs differentiated, intra-donor). *A*_*reg*_: PCI of log no1malized *F*_*reg*_ counts. *A*_*rand*_: PCI of 2000 random peaks matched for mean accessibility.

### Hostile controls

**Cl:** Depth+ cell cycle residualization. **C2:** Block permutation (N=5 per donor-state). **C3:** Feature randomization. **C4:** Label **shuffle—State-label shuffling within donors abolishes CE/CS differences (data not shown). CS:** Threshold sensitivity.

### Statistical testing

Wilcoxon signed-rank test (paired, donor-level). N=10 donors for CE/CS; N=12 for GO. Two-sided tests; *α* = 0.05.

## Data and Code Availability

Primary data: GEO accession GSE194122. Analysis code and outputs frozen with SHA256 checksums (manifest_files.tsv).

## Acknowledgments

We thank the authors of GSE194122 for public data access.

## Author Contributions

E.N.Q. conceived LCR-DE framework, designed metrics, supervised analysis. R.N.Q. implemented computations, generated figures, performed hostile controls. Both authors wrote the manuscript.

## Supplementary Figure Legends

**Supplementary Figure 1. Site Robustness: CE and CS patterns reproduce across collection sites**. Top panel: CE by collection site (site1-site4), stratified by state. Progenitor (blue) > Differentiated (coral) pattern evident across all sites. Bottom panel: CS by site. Differentiated > Progenitor pattern consistent. Inter-site variance reflects donor heterogeneity within sites, not batch effects. No site reverses pattern direction.

## Supplementary Tables

**Table Sl:** Donor × state cell counts. Complete cohort inventory.

**Table S2**: GO thresholds per donor. Donor-specific 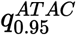 and 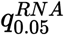 values (loglp units) computed on all cells before state subsetting.

**Table S3:** CE by donor × state with confidence intervals.

**Table S4:** CS by donor × state with component MI values.

**Table S5:** Permutation **null** statistics (N=5 permutations per donor-state).

**Table S6:** Hostile controls audit summary.

**Table S7:** Wilcoxon test statistics for CE and CS.

**Table S8:** Cell cycle scores and CE/CS before and after residualization.

**Table S9:** Wilcoxon test for cell cycle control (CE: p=0.64; CS: p=0.002).

**Table S10:** Site-stratified CE and CS summary statistics.

