## Supplementary material for "State-Dependent Regulatory Compression: Chromatin Geometry Gates Information Flow in Hematopoiesis": README_reproducibility.md

# Reproducibility: Paper 2 v2 (Cell Cycle & Robustness)

This package contains the hardened results for LCR-DE (Paper 2), addressing Hostile Reviewer concerns regarding Cell Cycle confounders and Site Robustness.

## Contents
- **figures/Figure_P2_4_CellCycle_Control.pdf**: Proof that CE/CS trends persist after regressing out S/G2M scores.
- **figures/Figure_P2_5_SiteRobustness.pdf**: Verification that effects are consistent across collection sites.
- **tables/**: Source data for all controls.

## Execution
Run `scripts/analysis_cellcycle_control.py` and `scripts/analysis_robustness.py` to regenerate.
Manifest verified via SHA256.
