## Supplementary material for "State-Dependent Regulatory Compression: Chromatin Geometry Gates Information Flow in Hematopoiesis": empirical_audit.md

# Empirical Audit Report

All figures and tables are derived exclusively from empirical measurements; no simulated or synthetic data were used.

## Verified Artifacts
- GO Thresholds: Derived intra-donor (Table S2)
- Cell Cycle: Scored via Tirosh gene set (Table S8)
- Feature Selection: Differential peaks (Empirical)
