## Supplementary material for "State-Dependent Regulatory Compression: Chromatin Geometry Gates Information Flow in Hematopoiesis": narrative_alignment.md

# Narrative Alignment Report

Paper 2 addresses mechanism, not repetition of the falsification in Paper 1.
- Falsification of Decoupling: Done (Paper 1).
- LCR-DE Mechanism: Done (Paper 2 Main Figs 1-4).
- CS Interpretation: 'Distributed Regulatory Redundancy' in Progenitors.
