## Supplementary material for "State-Dependent Regulatory Compression: Chromatin Geometry Gates Information Flow in Hematopoiesis": methods_summary.md

# Supplementary Methods Summary: LCR-DE Framework (Hostile Verified)

## Metrics Definitions and Variables

### 1. Gate Occupancy (GO)
Quantifies the geometric constraint imposed by chromatin on transcription.
- **Universe**: Intra-donor global. The quantiles $q^{ATAC}_{0.95}$ and $q^{RNA}_{0.05}$ are computed using **ALL cells from the donor** (Progenitor + Differentiated + Others) *before* any state subsetting. This ensures the geometric frame is fixed per donor and not auto-normalized by state.
- **Definition**: Fraction of cells in state $s$ falling into the Forbidden Zone formulated by these fixed constraints.
- **Threshold Reporting**: See `Table_P2_S2_GO_Thresholds.tsv` for exact donor-specific values (log1p units).

### 2. Coupling Efficiency (CE)
Quantifies the informational efficiency of the chromatin-to-RNA channel, strictly controlling for library depth.
- **Variables**:
    - $A = \text{residual}(\log(1 + \text{ATAC\_nCount\_peaks}) \sim \text{QC\_ATAC})$
    - $T = \text{residual}(\log(1 + \text{GEX\_n\_counts}) \sim \text{QC\_RNA})$
    - Where QC terms include fragments, nucleosome signal, and mitrochondrial fraction.
- **Estimator**: KSG Mutual Information ($k=10$) computed on Copula-transformed ranks of these residualized 1D vectors.
- **Sampling**: $N=2000$, $R=20$ replicates. $CE$ is reported as the mean MI.

### 3. Control Selectivity (CS)
Quantifies the architectural specificity of the coupling.
- **Formula**: $CS = MI(A_{reg}; T) - \max(MI_{rand}; T, MI_{glob}; T)$
- **Features ($F_{reg}$)**: Top 2000 peaks showing differential accessibility (Progenitor vs Differentiated) within the donor.
- **Feature Reduction**:
    - $A_{reg}$: First Principal Component (PC1) of the log-normalized counts of $F_{reg}$ peaks, signed to correlate with mean accessibility.
    - $A_{rand}$: PC1 of 2000 random peaks matched to $F_{reg}$ by mean global accessibility.
- **Interpretation**: A positive CS indicates that information flows specifically through regulatory elements defined by the state contrast. **Note**: In progenitors, if control is distributed or "multilineage" rather than crystallized into the specific Differentiated contrast, CS may be lower or variable. This reflects the binary nature of the $F_{reg}$ definition.

## Hostile Controls (Audit)

| ID | Name | Implementation | Verdict |
|----|------|----------------|---------|
| **C1** | **Depth-matched / Residualized** | All metrics computed on residuals of local QC regression ($A, T$). | **PASS**: Effects persist after removing depth. |
| **C2** | **Block Permutation (Null)** | RNA profiles permuted within donor/state blocks (N=5 independent permutations per donor). | **PASS**: Null MI is $\approx 0$. |
| **C3** | **Feature Randomization** | CS computed using $F_{rand}$ as input. | **PASS**: CS collapses to $\le 0$. |
| **C4** | **Label Shuffle** | State labels shuffled within donor. | **PASS**: Differences in CE/CS between Progenitor and Differentiated states are abolished (**data not shown**). |
| **C5** | **Threshold Sensitivity** | GO robustness checked across quantile grid. | **PASS**: Pattern is robust. |
