## Supplementary material for "State-Dependent Regulatory Compression: Chromatin Geometry Gates Information Flow in Hematopoiesis": README_supplementary.md

# Supplementary Materials: Paper 2 (LCR-DE)

## Supplementary Figures
- **Supplementary_Figure_1_SiteRobustness.pdf**: Verification of consistency across collection sites.

## Supplementary Tables
- Tables S1-S10 source data provided as TSV.
- **Table S2**: Explicit GO Thresholds per donor.
- **Table S8**: Cell Cycle scores and control metrics.
