## Supplementary figures and images for "State-Dependent Regulatory Compression: Chromatin Geometry Gates Information Flow in Hematopoiesis"

### Main_Figure_1_LCR_Framework.pdf

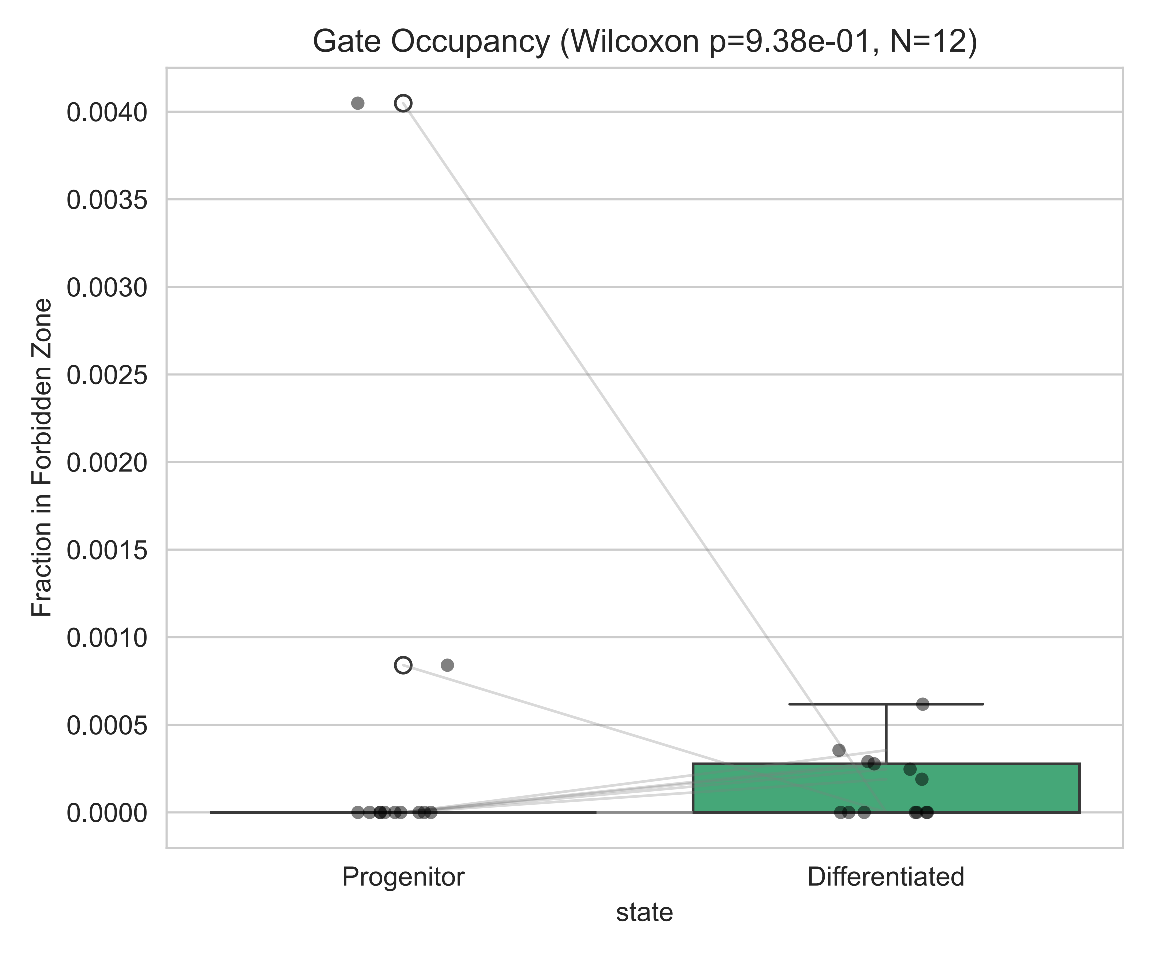

### Main_Figure_2_CE_by_State.pdf

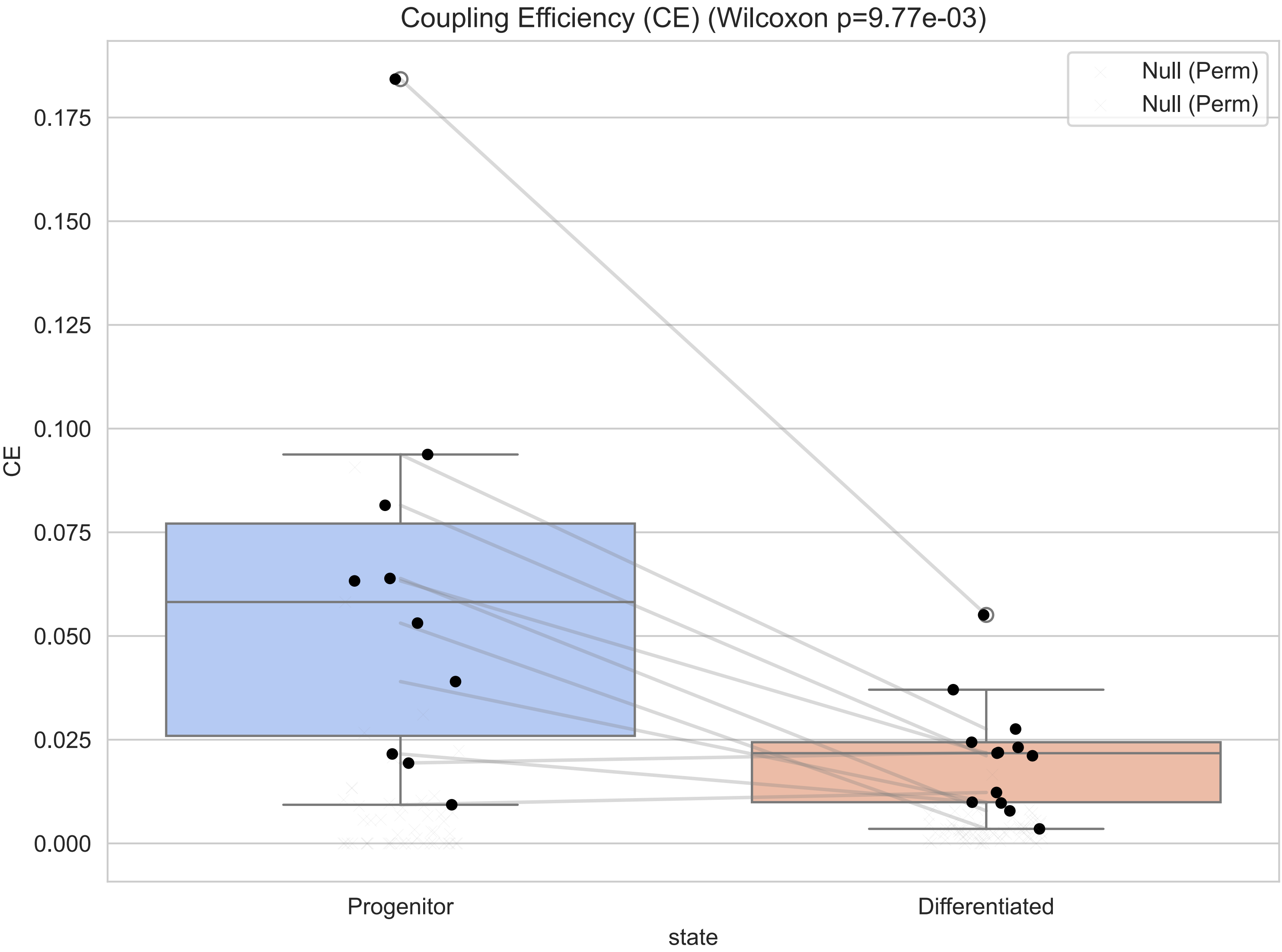

### Main_Figure_3_CS_Selectivity.pdf

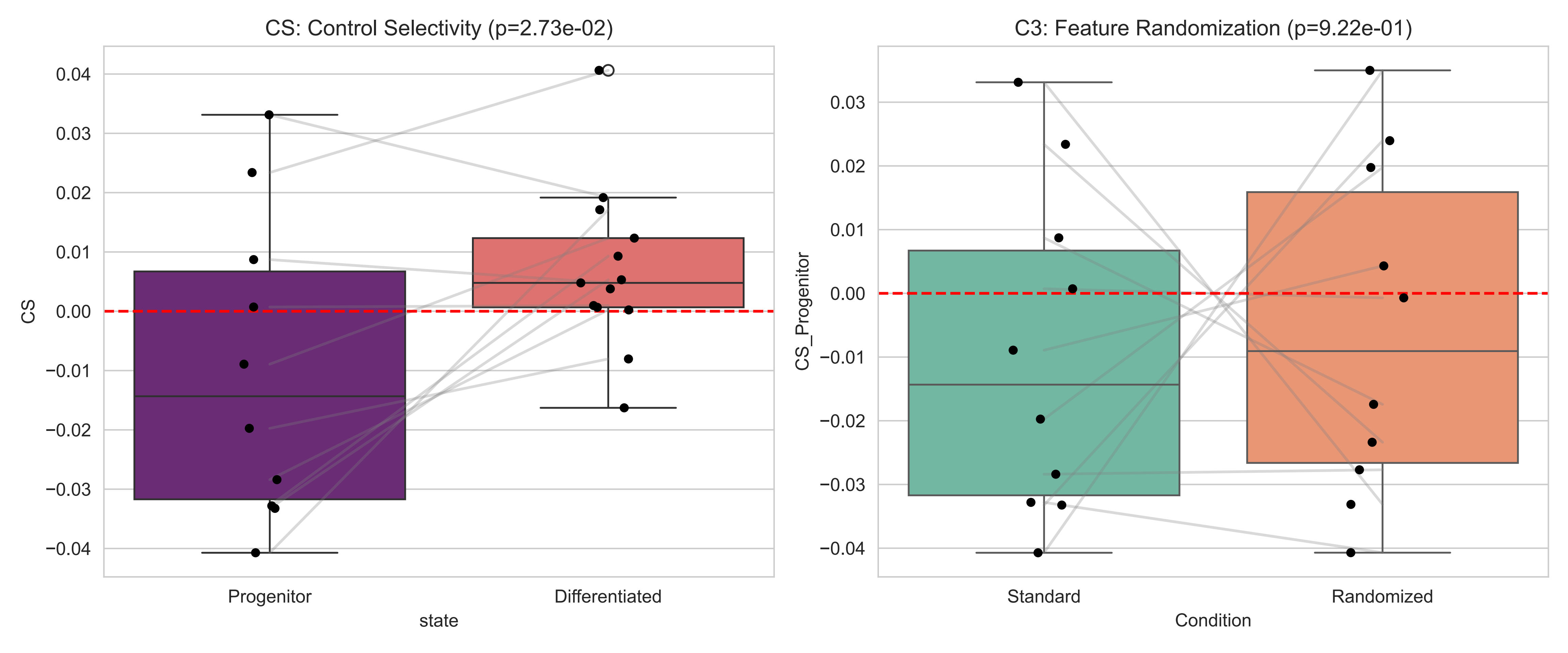

### Main_Figure_4_CellCycle_Control.pdf

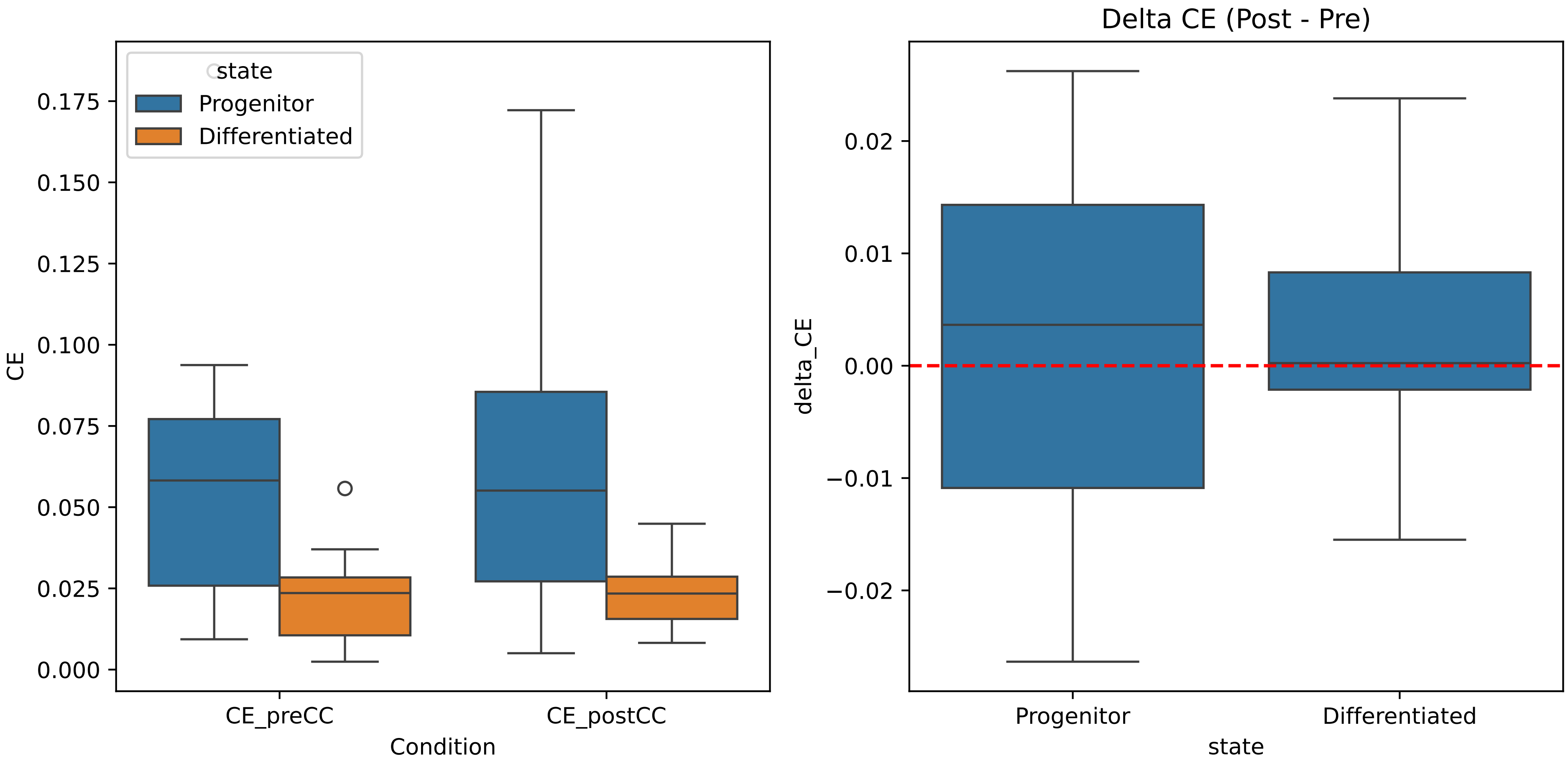

### Supplementary_Figure_1_SiteRobustness.pdf

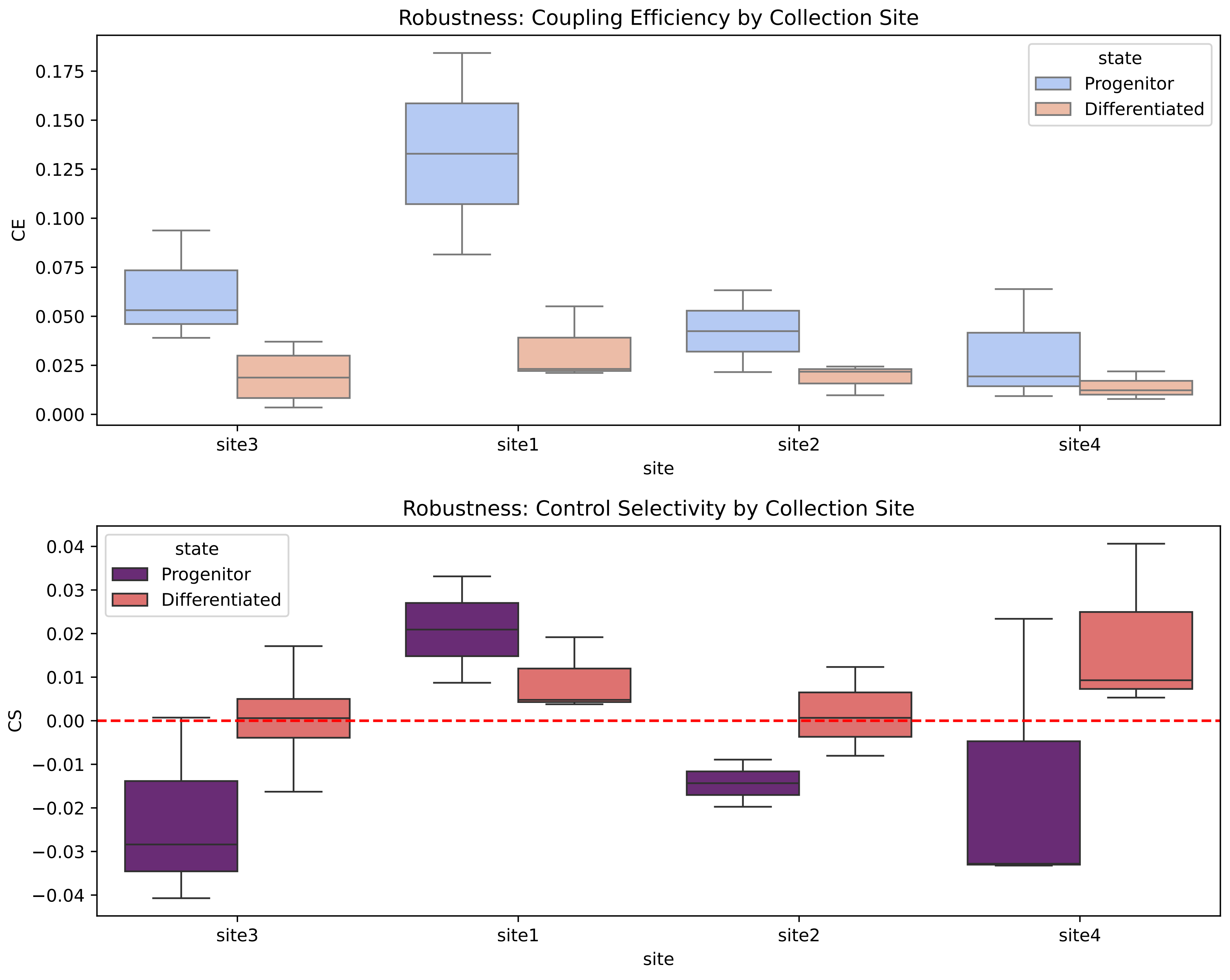
